# Dietary Indoles Modulate Gut Barrier Integrity via the AhR–IL-22 Axis in ART-Treated SIV Infection

**DOI:** 10.64898/2025.12.04.692392

**Authors:** Siva Thirugnanam, Alison R Van Zandt, Alexandra B McNally, Victoria A Hart, Isabelle Berthelot, Lara A Doyle-Meyers, David A Welsh, Andrew G MacLean, Namita Rout

## Abstract

HIV infection rapidly impairs the gastrointestinal (GI) barrier, contributing to persistent mucosal immune dysfunction, microbial translocation, and systemic inflammation despite antiretroviral therapy (ART). Using SIV-infected rhesus macaques on long-term ART, we investigated mechanisms underlying impairment in gut barrier-protective IL-17/IL-22 responses and the potential modulation of this pathway by dietary indoles. Longitudinal profiling of colonic epithelial and lamina propria cells revealed a selective loss of IL-17/IL-22–producing γδ T cells and type 3 innate lymphoid cells (ILC3s). This loss correlated with reduced expression of the transcription factors AhR and RORγt and was associated with elevated plasma markers of intestinal epithelial barrier disruption (IEBD), including intestinal fatty acid–binding protein (iFABP), zonulin, and LPS-binding protein (LBP). Targeting this transcriptional deficiency, dietary indole supplementation for one month restored colonic AhR⁺ IL-22–producing γδ T cells and RORγt⁺ ILC3s and Vδ1 T cells, and was associated with reduced iFABP and zonulin levels. Our findings indicate that disruption of the AhR–RORγt–IL-17/IL-22 axis is a key pathogenic mechanism underlying persistent IEBD in chronic SIV/HIV infection. Modulation of gut AhR signaling may represent a potential approach to reinforce mucosal barrier function and reduce chronic inflammation that persists in people living with HIV.

## Introduction

Chronic HIV/SIV infection is characterized by persistent immune activation and systemic inflammation, even with the successful suppression of viral replication through antiretroviral therapy (ART)^1, 2^. A key driver of this pathology is the damage to the gastrointestinal tract, leading to a phenomenon known as “leaky gut”^3^. This intestinal barrier dysfunction allows microbial products to translocate from the gut lumen into the bloodstream, fueling the chronic inflammation that contributes to incomplete immune recovery^3^ and non-AIDS comorbidities including cardiovascular, kidney, and liver diseases^4^. While ART is highly effective at controlling viremia and reducing some inflammatory markers, it often fails to fully restore the gut barrier and immune homeostasis^5, 6^, highlighting the need for novel therapeutic strategies that specifically target the mechanisms of chronic inflammation and intestinal dysfunction.

HIV and SIV preferentially infect intestinal CD4⁺ Th17 cells, whose loss is linked to epithelial barrier dysfunction (IEBD), microbial translocation, systemic inflammation, and disease progression^5, 7^. Chronic HIV also depletes protective Vδ2 γδ T cells while expanding inflammatory Vδ1 subsets, contributing to barrier damage^8^. Vδ2 cells transiently expand and produce IL-17/IL-22 during acute SIV, then decline, inverting the Vδ2:Vδ1 ratio^9^, suggesting a short-lived compensatory role. Elite and viral controllers maintain higher Vδ2 frequencies and IL-17 production than untreated or ART-treated people with HIV (PWH)^10, 11^. Group 3 innate lymphoid cells (ILC3s) similarly support barrier integrity^12, 13^ but decline during long-term ART^13, 14^, correlating inversely with plasma iFABP. The mechanisms underlying functional loss of Type 3 lymphocytes (Th17, γδ T, ILC3) in chronic treated infection, and whether they can be modulated to restore barrier function, remain unclear.

The functional integrity of Type 3 effector lymphocytes is dependent on key transcription factors, such as T-box protein expressed in T cells (T-bet), which drives Th1 cell polarization, and transcription of the retinoic acid-related orphan receptor γ isoform t (RORγt), which is essential for ILC3, γδ T, and Th17 cell development and function^15–17^. The Aryl Hydrocarbon Receptor (AhR), a ligand-activated transcription factor, also plays a significant role in regulating mucosal immunity and barrier function^16^. AhR binding is major inducer of transcription of RORγt, and activating AhR signaling via dietary or microbiota-derived ligands was shown to restore of intestinal barrier integrity and function via IL-22 and IL-17 secretion, strengthening of tight junctions, and induction of IL-10R^18^.

Despite their critical roles, the dynamics of Type 3 effector lymphocytes during acute and chronic SIV infection, and their response to long-term ART, remain incompletely characterized. Here, we define the compartmentalized transcriptional and functional profiles of circulating and colon-resident γδ T cells and ILC3s across the course of SIV infection and ART in rhesus macaques. We further evaluate the effects of dietary AhR ligand supplementation on gut barrier integrity and mucosal immune function. Our findings highlight a potential role for the AhR-RORγt-IL-22 axis in supporting gut barrier integrity and limiting chronic inflammation in ART-treated HIV/SIV infection.

## Results

### iFABP Recovery During ART Parallels Decline in SIV-Induced Proinflammatory Cytokines

Eleven rhesus macaques were infected with SIV and followed for over 6 months, with a subgroup of 5 receiving broccoli-based dietary AhR ligands to target gut inflammation (Fig. 1A). Plasma viremia peaked at 2 weeks (∼7.04 log₁₀ copies/mL) and stabilized at ∼2.96 log₁₀ copies/mL by 4 weeks (Fig. 1B). Daily ART (TDF, FTC, DTG) began at 6 weeks post-infection, achieving viral suppression below the assay LOD within 6 weeks. Plasma inflammatory and intestinal epithelial barrier dysfunction (IEBD) biomarkers were measured at baseline, acute infection, early ART (2-4 months of ART), and late chronic ART phases (>5-months of ART).

**Fig. 1.**
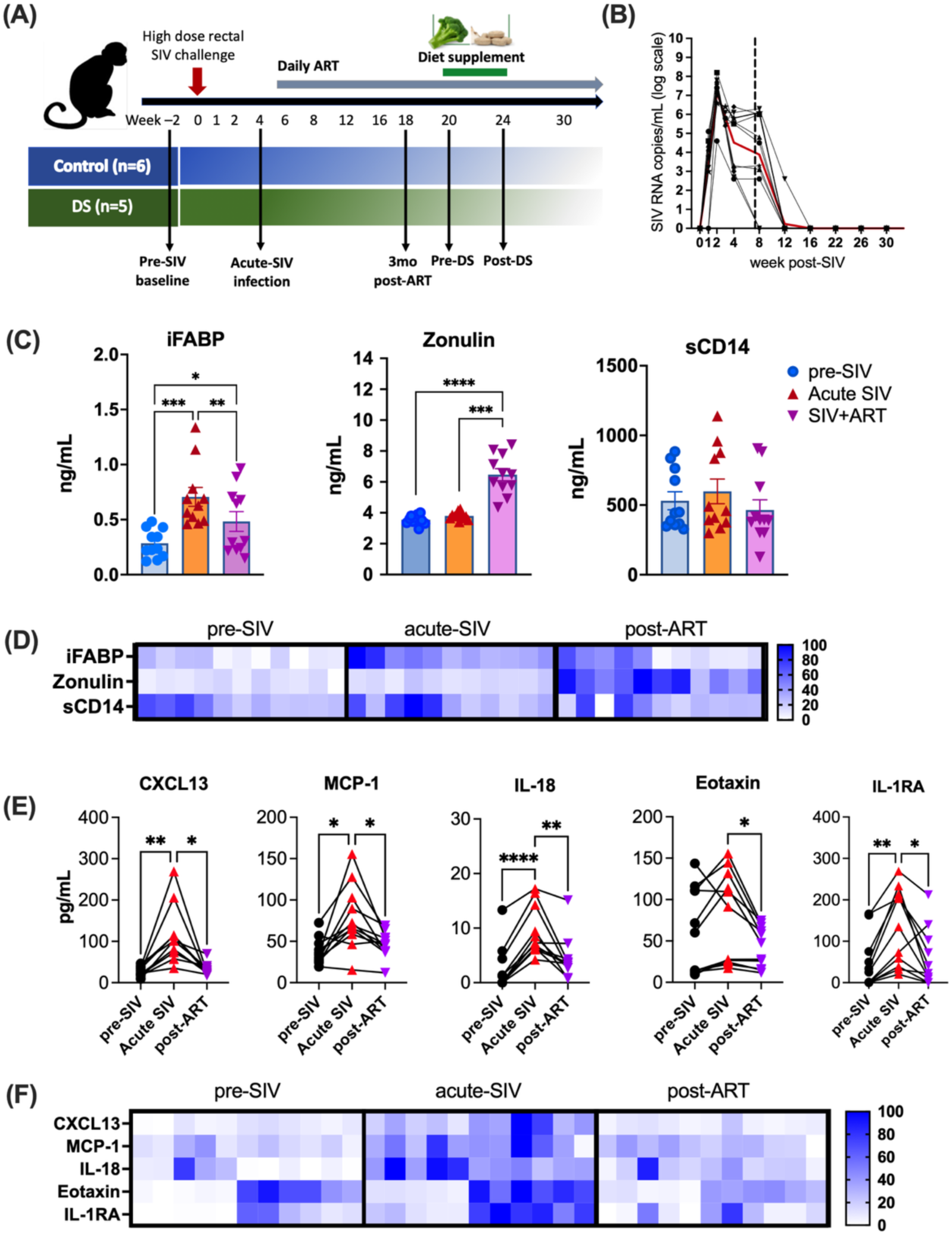
Plasma biomarkers of epithelial barrier disruption, microbial translocation, and inflammatory cytokines during SIV infection and ART. **(A)** Study design. SIV-infected rhesus macaques (n = 11) received daily ART (TDF, FTC, DTG). Baseline blood and gut biopsy samples were collected at week −2 and d0 SIV challenge, and ART was initiated at 6 weeks post-SIV infection. **(B)** Plasma SIV RNA levels over 38 weeks. Dashed line marks ART initiation; red line shows group average. **(C)** Plasma iFABP, Zonulin, and sCD14 concentrations at pre-SIV baseline, 4-weeks post-SIV infection (acute SIV), and 12-weeks post-ART. **(D)** Heatmap showing relative concentrations of iFABP, Zonulin, and sCD14 on a scale of 0-100 for each analyte. **(E)** Plasma CXCL13, MCP-1, IL-18, Eotaxin, and IL-1RA levels and **(F)** heatmap showing their relative concentrations at baseline, acute SIV, and post-ART time-points. Mean ± SEM; paired ANOVA (*p<0.05, **p<0.01, ***p<0.001, ****p<0.0001).

iFABP, a marker of enterocyte damage, rose significantly during acute infection (4 weeks, P = 0.019) and remained elevated despite ART, whereas zonulin increased later, 3 months post-ART (P = 0.015), suggesting early epithelial injury triggers tight junction disruption (Fig. 1C-D). sCD14, a marker of microbial translocation, remained unchanged. Circulating proinflammatory cytokines (CXCL13, IL-18, Eotaxin, IL-1RA) surged at 4 weeks post-SIV and partially declined with ART (Fig. 1E-F). These findings indicate that SIV-induced IEBD occurs early, with epithelial damage leading to zonulin-mediated tight junction disruption and inflammation, and although ART partially normalizes iFABP and reduces inflammation, zonulin remains elevated beyond baseline, likely contributing to persistent microbial translocation and inflammation despite effective viral suppression.

### Depletion of Circulating Vδ2 T Cells and ILC3s During Acute SIV Infection Is Associated with Peak Viremia and Elevated α4β7 Expression

Extensive HIV/SIV replication in gut-associated lymphoid tissue (GALT) depletes CCR5⁺ and Th17 CD4⁺ T cells, disrupting the epithelial barrier and promoting microbial translocation. Compensatory mucosal populations, including γδ T cells and ILC3, support barrier integrity via IL-17 and IL-22 production^9, 13^. We tracked these populations during acute SIV infection and ART. γδ T cells were identified as CD3⁺ Vδ1 or Vδ2 TCR⁺ cells, and ILC3 as lineage⁻CD8α⁻CD127⁺CD161⁺ cells (Fig. 2A). Blood CD4⁺ T cells declined at 4 weeks post-SIV but recovered by 3 months post-ART (Fig. 2B). Vδ2 T cells mirrored this pattern, showing an early rise at 1-week post-SIV before returning to baseline, reflecting a proliferative response (Fig. 2C) corroborating our earlier findings^9^. On the other hand, a rapid and progressive decline in ILC3 beginning during peak viremia was observed (Fig. 2D). The kinetics of the decline in peripheral Vδ2 T cells and ILC3, though divergent at 1-week post-SIV, was consistent between 1-4 weeks of SIV infection in displaying an inverse relationship with plasma viral loads (Fig. 2E-F). Further, Vδ1 T, Vδ2 T, and ILC3 exhibited increased α4β7 expression, indicating enhanced gut homing during viral replication (Fig. 2G).

**Fig. 2.**
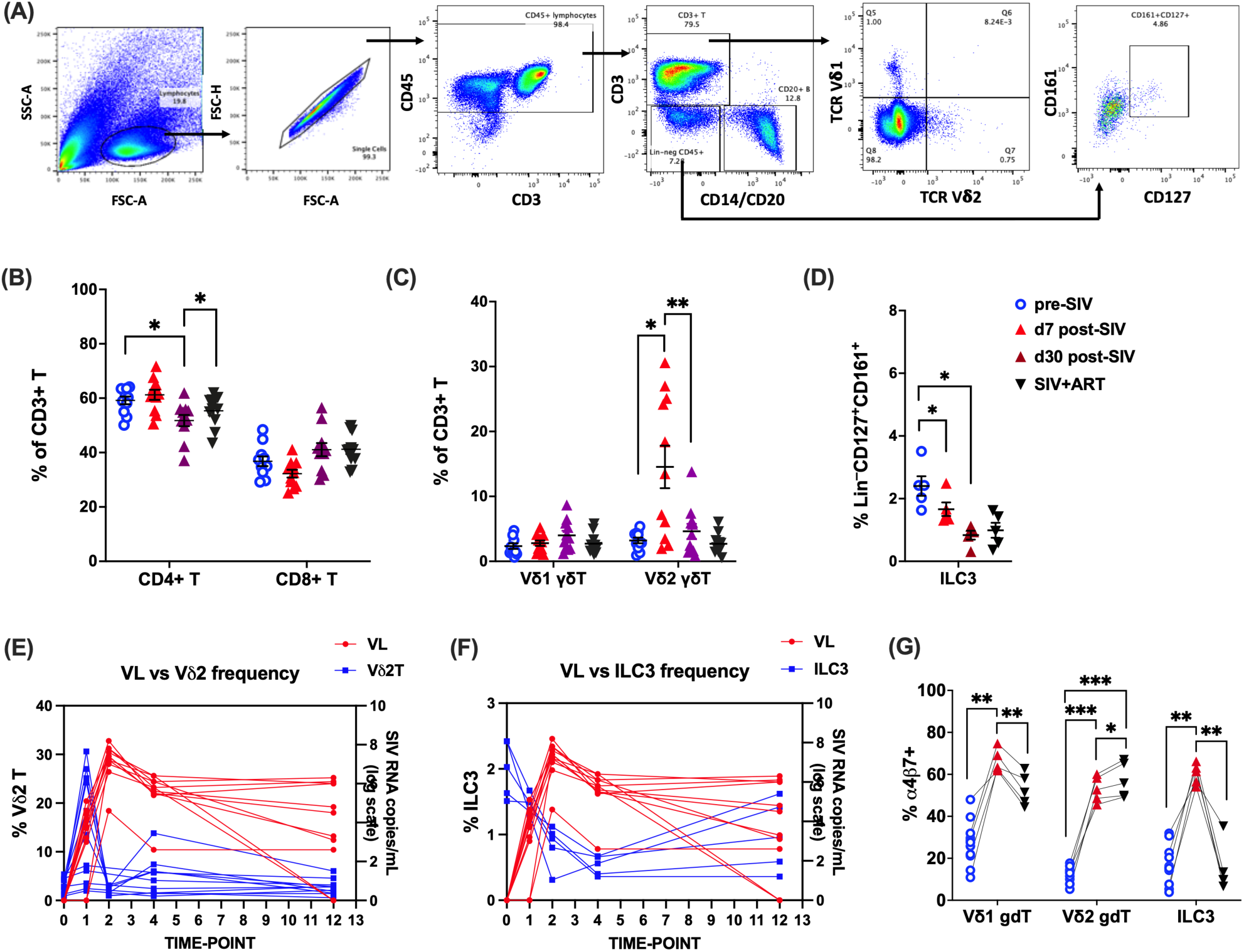
Temporal dynamics of Vδ2 T cell and ILC3 depletion during acute SIV infection and their relationship to peak viremia. **(A)** Representative gating schematic for Vδ1 T, Vδ2 T, and ILC3 in PBMC. **(B)** Frequencies of CD4+ and CD8+ T cells, **(C)** Vδ1 and Vδ2 γδT cells, and **(D)** ILC3 in peripheral blood of study animals at baseline (2 weeks pre-SIV infection), acute SIV infection (7d and 1-month post-SIV), and 12-week post-ART. Circulating immune cell frequencies (blue, left y-axis) and SIV viral load in log scale (red, right y-axis) at each sampling time point for **(E)** Vδ2 T cells and **(F)** ILC3. **(G)** Frequencies of PBMC Vδ1 T, Vδ2 T, and ILC3 expressing α4β7 at indicated time points. Mean ± SEM; paired ANOVA (*p<0.05, **p<0.01, ***p<0.001, ****p<0.0001).

### SIV-Driven Th1 Polarization Persists in Circulating γδ T Cells and ILC3s During ART, Marked by T-bet Upregulation and RORγt/AhR Downregulation

The shift in Th1/Th17 balance in mucosal tissues is a well-documented hallmark of HIV/SIV infections^19, 20^. To examine changes in functional immunophenotype of circulating ILC3 and γδ T cells, changes in their transcription factor profiles were evaluated at baseline (Fig. 3A-C), following SIV infection and during the immune reconstitution phase of ART (Fig. 3D-F). At baseline, γδ T cells and ILC3 exhibited higher frequencies of cells expressing the Th17-associated transcription factors AhR and RORγt compared to those expressing the Th1-associated factor T-bet (Fig. 3B). Notably, both γδ T cells and ILC3s also showed significantly higher mean fluorescence intensity (MFI) of RORγt relative to T-bet, indicating a predominant Th17-type functional phenotype (Fig. 3C). Despite the higher frequencies of AhR-positive γδ T cells and ILC3s, the per-cell expression of AhR was relatively low, as reflected by low MFI values. Among the subpopulations—Vδ1 T cells, Vδ2 T cells, and ILC3s—ILC3s demonstrated significantly higher frequencies of AhR expression and greater RORγt MFI, suggesting a stronger skewing toward Th17-type functionality (Fig. 3B-C).

**Fig. 3.**
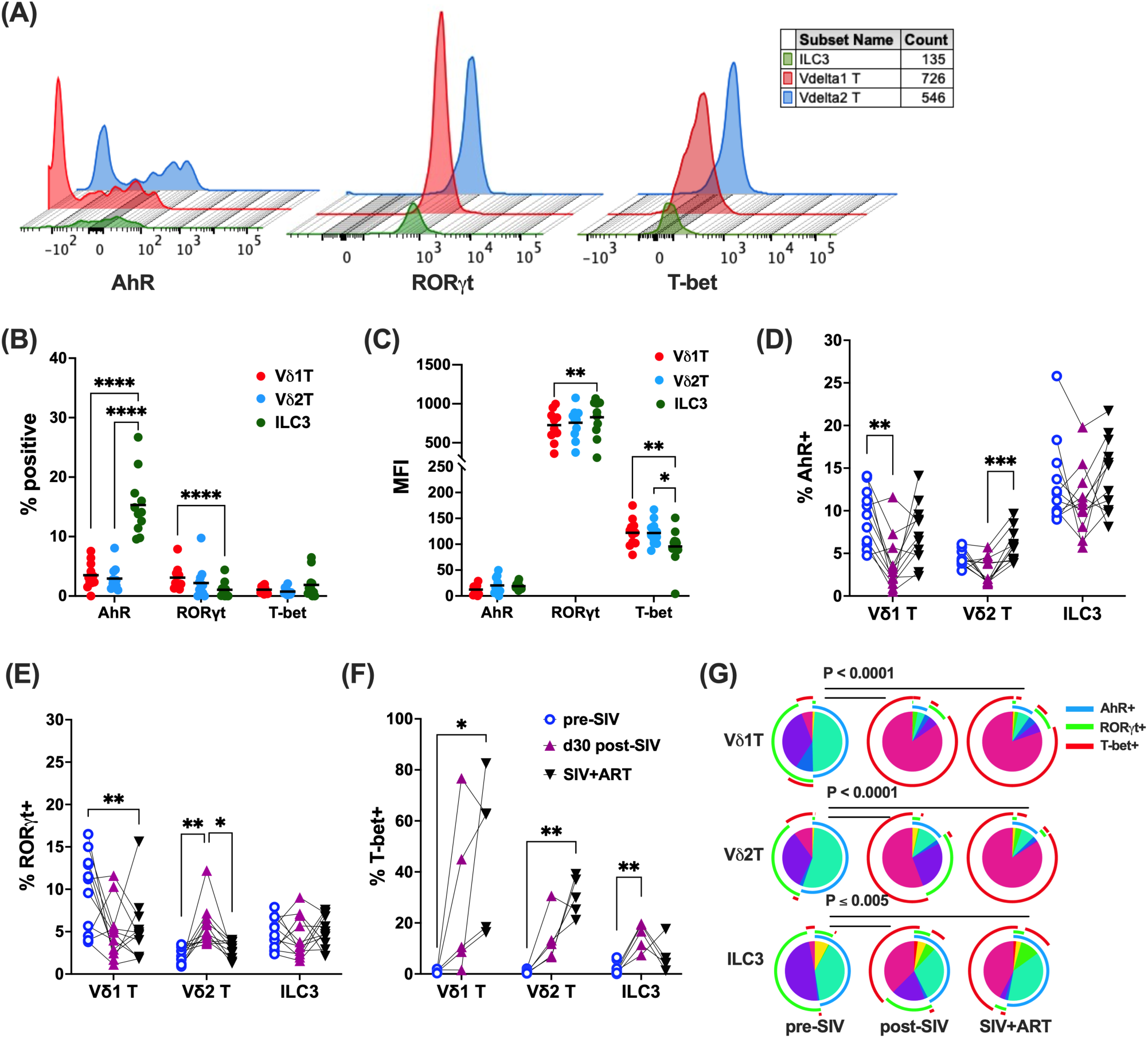
Transcription factor expression in circulating Type-3 immune cells during SIV infection and ART. **(A)** Histogram plot showing expression of the transcription factors AhR, RORγT, and T-bet in Vδ1 T, Vδ2 T, and ILC3 cells. **(B)** Frequencies of AhR, RORγT, and T-bet expression in PBMC Vδ1 T, Vδ2 T, and ILC3 subsets. **(C)** Mean fluorescence intensity (MFI) of AhR, RORγT, and T-bet on the Vδ1 T, Vδ2 T, and ILC3 populations in PBMC. Longitudinal changes in the frequencies of AhR+ **(D),** RORγT+ **(E),** and T-bet+ (F) Vδ1 T, Vδ2 T, and ILC3 populations evaluated at baseline (pre-SIV), acute SIV infection (d30 post-SIV), and 12-weeks post-ART (SIV+ART). Mean ± SEM; paired ANOVA (*p<0.05, **p<0.01, ***p<0.001, ****p<0.0001). **(G)** Pie charts depicting the relative expression of AhR, RORγT, and T-bet expression by Vδ1 T, Vδ2 T, and ILC3 populations from pre-SIV baseline through SIV infection and ART. Pie arcs represent expression of individual transcription factors and pie slices represent the number of co-expressed transcription factors. Significant differences between each time-point for the overall phenotype were evaluated by SPICE permutation test.

Following SIV infection, AhR⁺ Vδ1 T cells transiently declined but normalized with ART (Fig. 3D). Vδ2 T cells, by contrast, showed no significant change in AhR expression during acute infection but exhibited increased AhR expression during ART, indicating enhanced AhR signaling in the setting of viral control. Notably, Vδ2 T cells also showed an early increase in RORγt expression post-infection, which returned to baseline after two months on ART, coinciding with an upregulation of T-bet (Fig. 3E-F). These findings suggest that although ART restores the frequency of certain Th17-associated transcriptional profiles such as AhR^+^ Vδ2 T cells and RORγt^+^ Vδ1 T cells but may also promote a shift in Type-3 immune cells from a Th17-like phenotype (AhR^+^/RORγt^+^) toward a Th1-like phenotype (T-bet^+^) during acute infection and viral suppression (Fig. 3G). This dynamic may reflect immune reprogramming associated with treated HIV infection, where ART partially restores immune homeostasis but may also skew innate-like T cell subsets toward Th1-type responses.

### Compartmentalized γδ T Cell and ILC3 Dynamics in Colonic Epithelium versus Lamina Propria During SIV and ART

To further understand the gut-specific impacts of SIV infection and the influence of viral suppression with ART on γδ T cells and ILC3, we next examined their frequencies and immunophenotype in colonic lamina propria (LPL) and intraepithelial (IEL) compartments. As expected, a significant loss of CD4 T cells was observed in colonic lamina propria lymphocytes (LPL; P<0.0001) and intraepithelial lymphocytes (IEL; P<0.01) by 6 weeks post-SIV infection resulting in near-complete depletion of CD4⁺ T cells in this gut compartment around the time of set-point viremia (Fig. 4A, D) in this section of gut around the time of set-point viremia. Three months of daily ART partially restored CD4⁺ T cells in both IEL and LPL compartments (P<0.05; Fig. 4A, D). However, CD4⁺ T cell frequencies in the LPL remained significantly below baseline levels (p = 0.0058), indicating that viral suppression by ART promotes differential recovery of CD4⁺ T cells in the colonic lamina propria versus the epithelium.

**Fig. 4.**
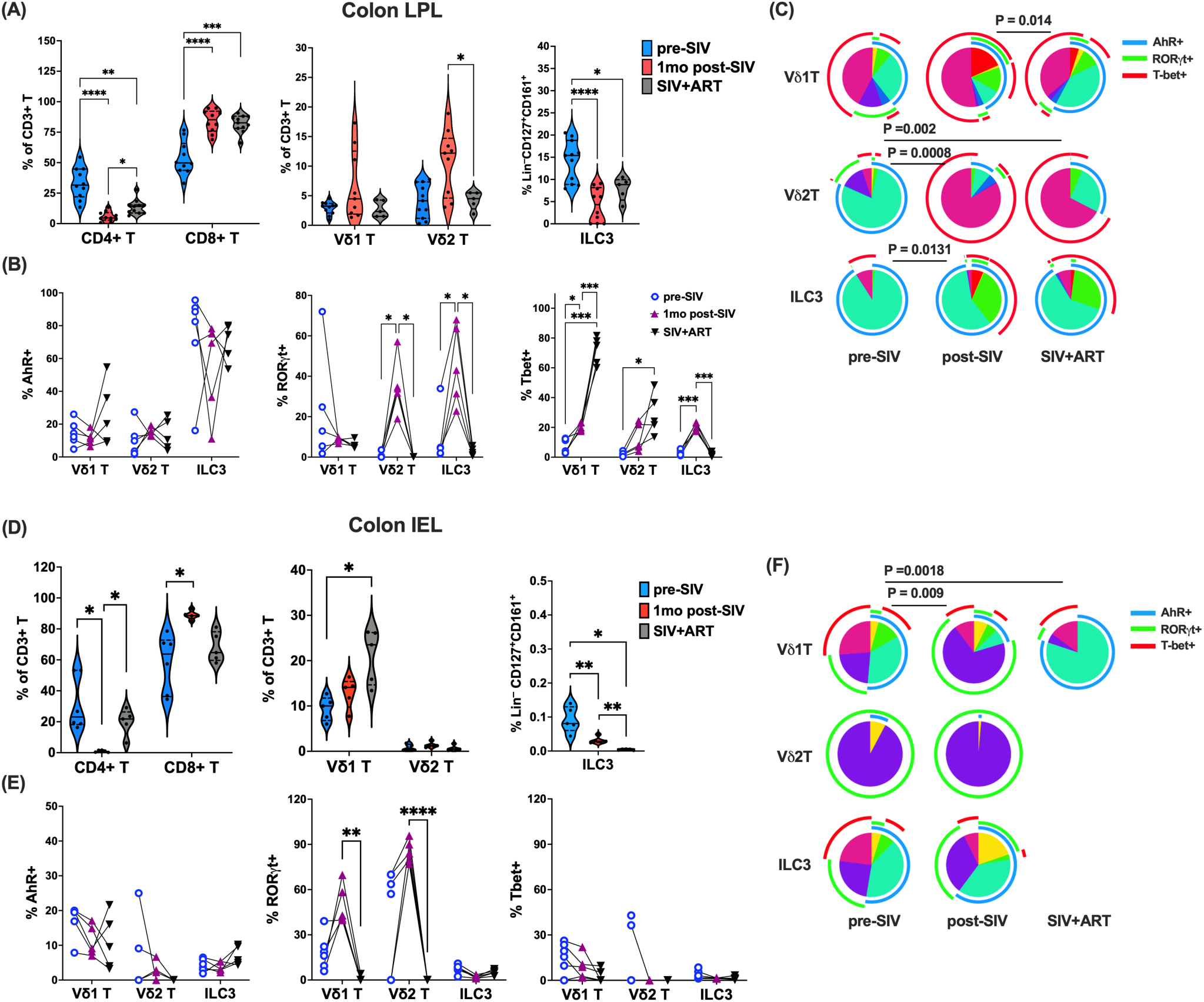
Frequencies of colonic mucosal γδ T cells and ILC3 and expression of Th1/Th17-associated transcription factors during acute SIV infection and ART. **(A)** Longitudinal frequencies of CD4^+^T, CD8^+^ T, Vδ1 T, Vδ2 T, and ILC3 populations in the LPL from colon biopsies at indicated time-points. **(B)** AhR, RORγT, and T-bet expression by colonic LPL Vδ1 T, Vδ2 T, and ILC3 populations. **(C)** Pie charts depicting the relative expression of AhR, RORγT, and T-bet expression by Vδ1 T, Vδ2 T, and ILC3 populations in colonic LPL isolated from pre-SIV baseline through SIV infection and ART. **(D)** Longitudinal frequencies of CD4^+^T, CD8^+^ T, Vδ1 T, Vδ2 T, and ILC3 populations in colonic IEL at indicated time-points. **(E)** AhR, RORγT, and T-bet expression by colonic IEL Vδ1 T, Vδ2 T, and ILC3 populations. **(F)** Pie charts depicting the relative expression of AhR, RORγT, and T-bet expression by Vδ1 T, Vδ2 T, and ILC3 populations in colonic IEL fraction from pre-SIV baseline through SIV infection and ART. Pie arcs represent expression of individual transcription factors and pie slices represent the number of co-expressed transcription factors. The rare frequencies of IEL Vδ2 T, and ILC3 populations precluded a meaningful analysis of their transcription factor profiles at the SIV+ART time-point. Mean ± SEM; paired ANOVA (*p<0.05, **p<0.01, ***p<0.001, ****p<0.0001).

In LPL, Vδ2 T cells increased during acute infection and returned to baseline by 3 months post-ART (P = 0.017), while Vδ1 T cells remained stable (Fig. 4A). However, ILC3s mirrored CD4⁺ T cell kinetics, sharply declining at 1 month post-infection (P<0.0001) with partial ART-mediated restoration (Fig. 4A). Notably, a transient Th17-type skewing, reflected by elevated RORγt, occurred in Vδ2 T cells and ILC3s during acute infection (P ≤ 0.012), whereas AhR remained unchanged across all three cell populations during acute SIV infection and after ART (Fig. 4B). Progressive T-bet upregulation was noted in γδ T cells, most pronounced in Vδ1 (P = 0.0004), while T-bet⁺ ILC3s increased transiently during acute infection before returning to baseline (Fig. 4B). These findings suggest that during acute SIV infection, Vδ2 T cells and ILC3s undergo transient Th17-type skewing in response to viral replication and CD4⁺ Th17 cell depletion in the gut (Fig. 4C). Furthermore, the sustained T-bet upregulation in Vδ1 T cells, despite effective viral suppression with ART, points to a potential role for this subset in maintaining mucosal inflammation during chronic infection (Fig. 4C).

The colonic intraepithelial lymphocyte (IEL) compartment exhibited a near-complete depletion of CD4⁺ T cells following SIV infection, accompanied by a reciprocal increase in CD8⁺ T cell frequencies (Fig. 4D), highlighting a major shift in T cell composition at the mucosal barrier. Vδ2 T cells remained nearly undetectable at all timepoints, consistent with prior findings that Vδ1 T cells predominate among γδ T cells in the gut epithelium^21, 22^. Similarly, lineage-negative CD127⁺ ILC3-like cells, relatively rare subset in the IEL^23^, showed a progressive and sustained decline post-infection that was not reversed by ART (Fig. 4D), suggesting poor restoration of this critical innate population in the epithelial niche. Interestingly, IEL-resident Vδ1 T cells increased significantly by 3 months post-SIV+ART (P = 0.046), pointing to a selective expansion or enhanced retention of this subset in the epithelial compartment (Fig. 4D). Functionally, RORγt-expressing Vδ1 T cells were significantly elevated during acute infection (P = 0.0025), mirroring the Th17-like skewing observed in Vδ2 T cells in the LPL (Fig. 4E-F). However, this increase in RORγt expression was transient and declined to below baseline levels by 3 months post-ART (Fig. 4E). The low abundance of ILC3s precluded a meaningful analysis of their transcription factor profiles in the IEL. Thus, compartmentalized γδ T Cell and TCR-negative innate cell responses were observed in the colonic lamina propria and epithelium during SIV infection.

### Th1/Th17 Cytokine Dysregulation and Transcription Factor Correlates in Colonic γδ T Cells During ART-Treated SIV Infection

To assess how RORγt, AhR, and T-bet expression affects gut γδ T cell and ILC3 function, we measured cytokine production in colonic LPLs after ex vivo mitogen stimulation. Consistent with our earlier findings^9^, ex vivo production of Th17-type cytokines, IL-17A and IL-22, was markedly impaired during SIV infection despite ART, whereas IFN-γ production in mitogen-stimulated cells remained largely intact, except for a significant decline in CD4^+^ T cells (Supplementary Fig. 1). In colonic LPLs, IL-22 was significantly reduced in CD4⁺ (P = 0.0002) and Vδ2 T cells (P = 0.002), with a trend in ILC3s (P = 0.059), and IL-17 was decreased in CD4⁺ (P = 0.005) and Vδ2 T cells (P = 0.007) (Fig. 5A-B).

**Fig. 5.**
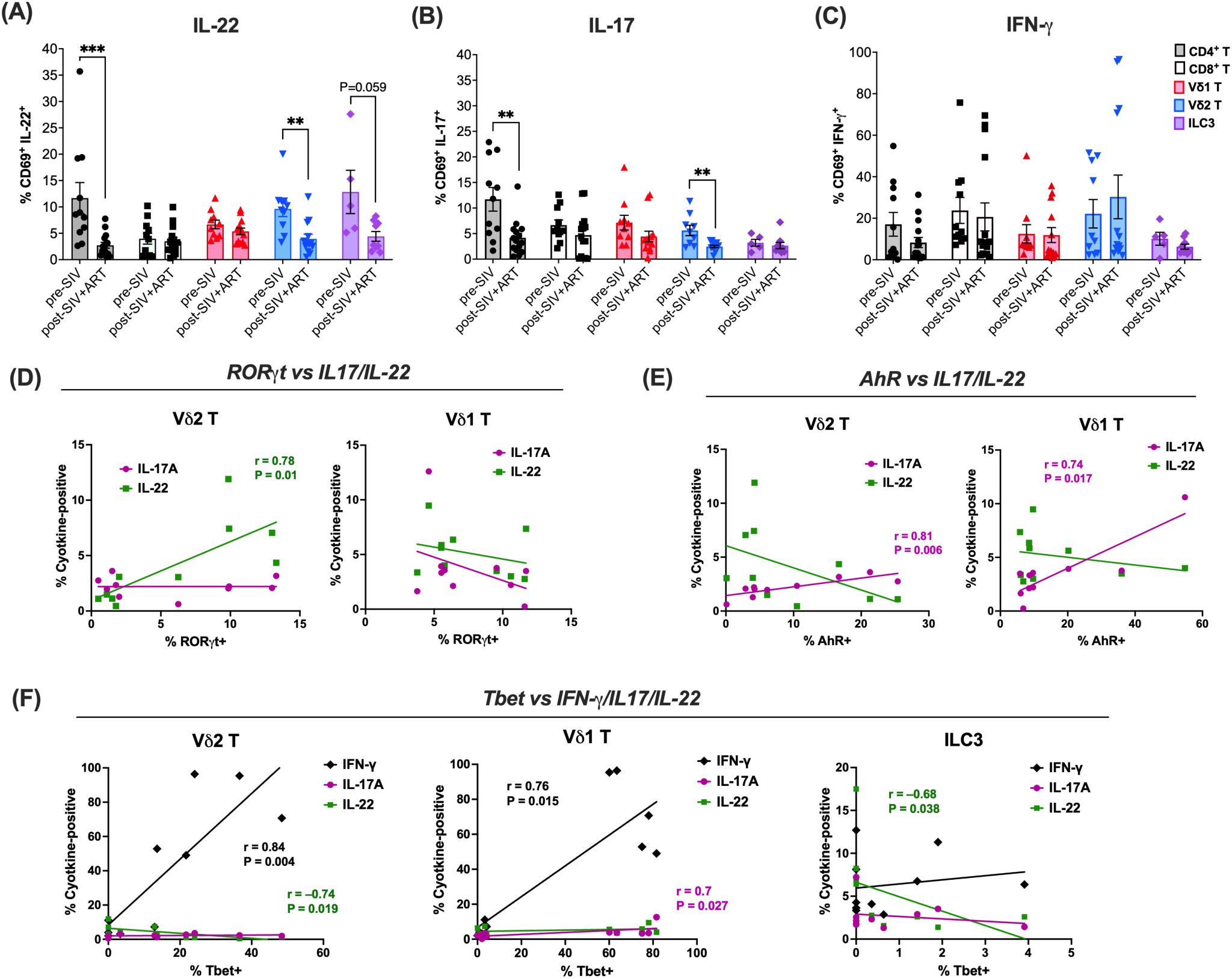
Cytokine production by colonic mucosal T cells and ILC3 during ART-suppressed SIV infection and correlation with transcription factor expression. Freshly isolated colonic lamina propria lymphocytes were evaluated for cytokine production following *ex vivo* stimulation with PMA/Ca Ionomycin. Frequencies of cells expressing IL-22 **(A)**, IL-17A **(B)**, and IFN-γ **(C)** were determined by intracellular flow cytometry and analysis of γδ T cells, CD4⁺ T cells, CD8⁺ T cells, and ILC3s. Data represent mean ± SEM from 5-15 animals per group. Expression of transcription factors was measured in parallel and correlated with cytokine output. Correlations of RORγt **(D)**, and AhR **(E)** expression with IL-17A and IL-22 cytokines produced by colonic γδ T cell subsets. Correlations of T-bet **(F)** with intracellular IFN-γ, IL-17A, and IL-22 cytokine expression by colonic γδ T cells and ILC3s. Mean ± SEM; Mann-Whitney test (*p<0.05, **p<0.01, ***p<0.001). Spearman correlation analysis was performed to assess the relationship between IL-17A and IL-22 and RORγt and AhR expression.

Notably, IL-22 expression in ex vivo stimulated Vδ2 T cells correlated positively with RORγt expression (P = 0.01; Fig. 5D), whereas IL-17 production was associated with AhR expression in both Vδ1 (P = 0.017) and Vδ2 T cells (P = 0.007; Fig. 5E), highlighting distinct transcriptional dependencies for these cytokines. As expected, T-bet expression strongly correlated with IFN-γ production in colonic γδ T cells (P = 0.004 in Vδ2; P = 0.015 in Vδ1; Fig. 5F). Strikingly, T-bet also negatively correlated with IL-22 in Vδ2 T cells and ILC3s (P = 0.019 and 0.038, respectively), consistent with the skewing toward an IFN-γ–dominant phenotype observed with increased T-bet and reduced RORγt in colonic LPLs (Fig. 4C). Interestingly, IL-17 production in Vδ1 T cells correlated positively with T-bet, suggesting that IFN-γ/IL-17 co-producing Vδ1 T cells are regulated via integrated T-bet and AhR signaling. No significant correlation was observed between AhR or RORγt expression and cytokine production in ILC3s (Supplementary Fig. 1). In summary, chronic SIV+ART infection reprograms mucosal γδ T cells and ILC3s toward a T-bet–driven, IFN-γ–biased phenotype with diminished IL-17 and IL-22 production.

### In Vitro Effects of AhR Ligands on Mitigating HIV-Induced Damage to Colonic Epithelial Monolayers

HIV proteins, including Tat, can disrupt intestinal epithelial integrity independent of viral replication^24^. We treated differentiated human colonic epithelial Caco-2 monolayers with recombinant SIV Tat protein as in vitro model. Tat-exposure resulted in significantly reduced cell viability comparable to 5–10% ethanol (Supplementary figure 2), and decreased AhR expression (Fig. 6A–B), suggesting that Tat-mediated damage may occur, at least in part, via AhR suppression. Immunofluorescence staining for the tight junction protein ZO-1 revealed disrupted junctional morphology in Tat-treated Caco-2 monolayers, characterized by diminished ZO-1 signal intensity and visible intercellular gaps (Fig. 6D; red arrows, inset B), indicative of compromised barrier integrity. Co-treatment with AhR ligands 6-formylindolo[3,2-b] carbazole (FICZ) and indole-3-carbinol (I3C) preserved ZO-1 organization and restored the characteristic “zigzag” junctional pattern associated with well-polarized, mature epithelial monolayers (Fig. 6C; insets C–D), with FICZ showing a stronger effect. Real-time cell analysis confirmed that Tat impaired barrier function (decreased impedance/Cell Index), which was significantly restored by FICZ and I3C, particularly FICZ (P < 0.0001), as evidenced by both real-time recovery kinetics and area under the curve (AUC) analysis (Fig. 6D). Because cruciferous vegetables such as broccoli contain indole glucosinolates hydrolyzed into AhR agonists like indolocarbazole (ICZ), we tested whether broccoli extract could activate AhR signaling. In the DR-EcoScreen luciferase assay, broccoli extract induced significant AhR activation in murine hepatoma cells, whereas blueberry extract, despite its polyphenol content, did not (Fig. 6E), demonstrating ligand-specific activation of AhR by dietary indoles.

**Fig. 6.**
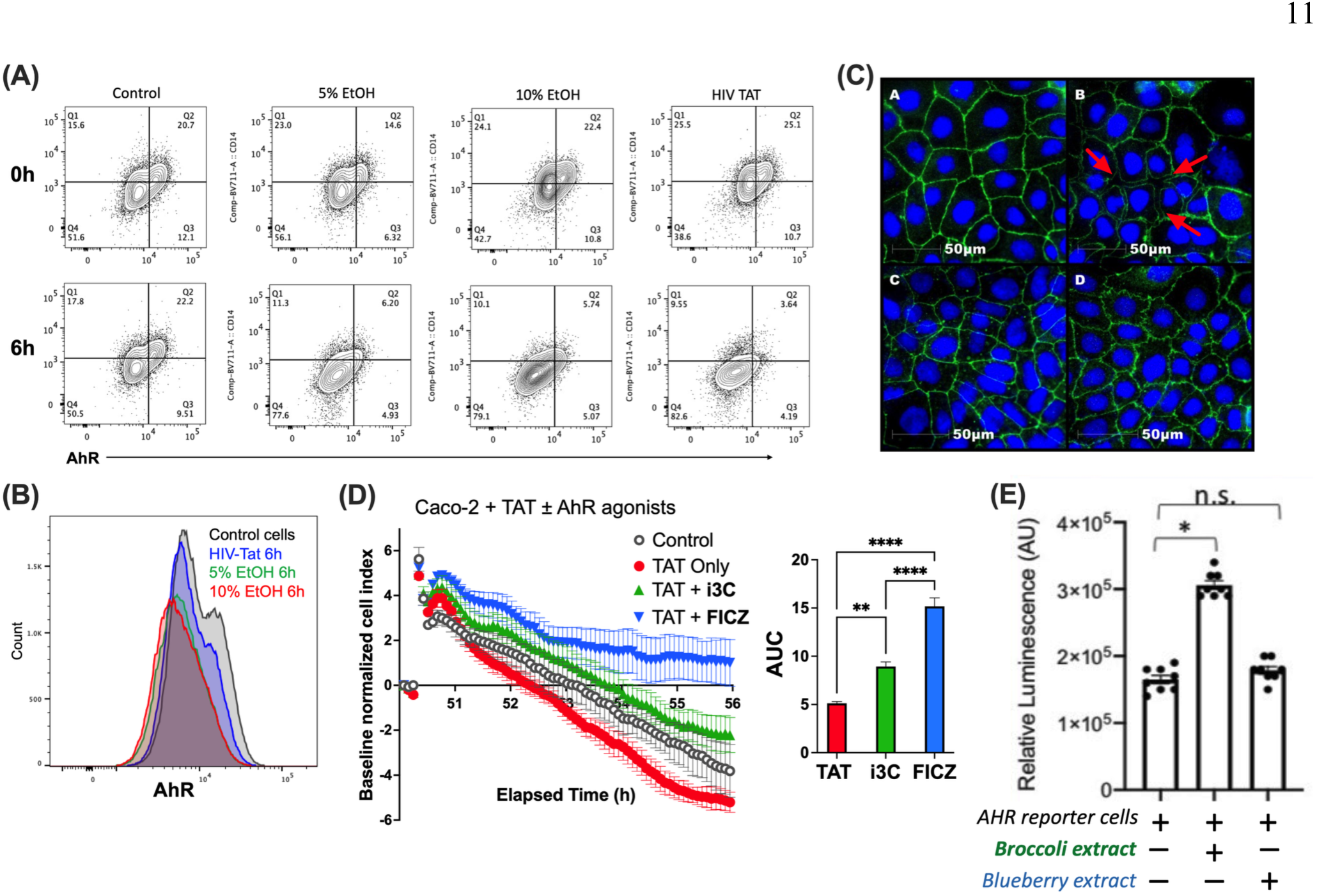
In vitro epithelial monolayer disruption and AhR downregulation by HIV Tat is reversed by ligand-specific activation through indoles. **(A)** Flow cytometry contour plots showing AhR expression in Caco-2 cells at 0-hour (top panel) and 6 hours (bottom panel) in control media or media supplemented with 5% EtOH, 10% EtOH, and 3μM HIV Tat. **(B)** Representative histogram showing AhR downregulation following 6-hour exposure to EtOH or Tat-containing media in comparison to control media. **(C)** Fluorescent staining of ZO-1 in untreated Caco-2 cells (inset A), and cells treated with Tat (inset B), TaT+i3C (inset C), and TaT+FICZ (inset D). **(D)** Normalized cell index of adherent Caco-2 cells evaluated by RTCA after incubation with Tat only (red symbols), Tat+i3C (green symbols) or TaT+FICZ (blue symbols) in comparison to control media (grey symbols). Each data point signifies the average ± standard deviation of triplicates. Area under curve (AUC) analysis of adjacent cell index data were plotted as mean ± SEM (**p<0.005, ****p<0.0001). **(E)** AHR activity evaluated by luciferase fluorescence quantification after 6-hour treatment of DR-EcoScreen luciferase reporter cells with either broccoli extract or blueberry extracts.

### Broccoli-Based AhR Ligand Supplementation Enhances Epithelial Barrier Integrity, Vδ2 T Cell Function, and Mucosal Immune Subset Distribution During Chronic SIV Infection and ART

We next assessed the effects of short-term broccoli-based dietary supplementation on AhR signaling and intestinal inflammation in chronically SIV-infected macaques on long-term ART. At 5 months post-SIV+ART, animals received the broccoli sprout extract Avmacol (950 mg/day sulforaphane) for 2 weeks, escalating to 1900 mg/day for 2 more weeks. Plasma biomarkers showed reduced iFABP by 2 weeks and LBP by 4 weeks (Fig. 7A-B), indicating improved barrier integrity and decreased microbial translocation. sCD14 decreased at 2 weeks, though not sustained in 2/5 animals (Fig. 7C). Concomitantly, dietary supplementation enhanced IL-17 responses in circulating Vδ2 T cells, as demonstrated by an increase in IL-17 spot-forming cells following HMBPP stimulation of PBMCs (Fig. 7D). Furthermore, at 4 weeks post-dietary supplementation (DS), there was a coordinated trend toward reduced systemic inflammation, including TNF-α, chemokines MCP-1 and CXCL13, and Eotaxin, suggesting a broad systemic anti-inflammatory effect. IL-1RA also declined, reflecting dampened negative feedback, while SDF-1α, FGF-2, and VEGF-D levels decreased, indicating reduced inflammatory cell recruitment and pro-angiogenic signaling (Supplementary Fig. 3).

**Fig. 7.**
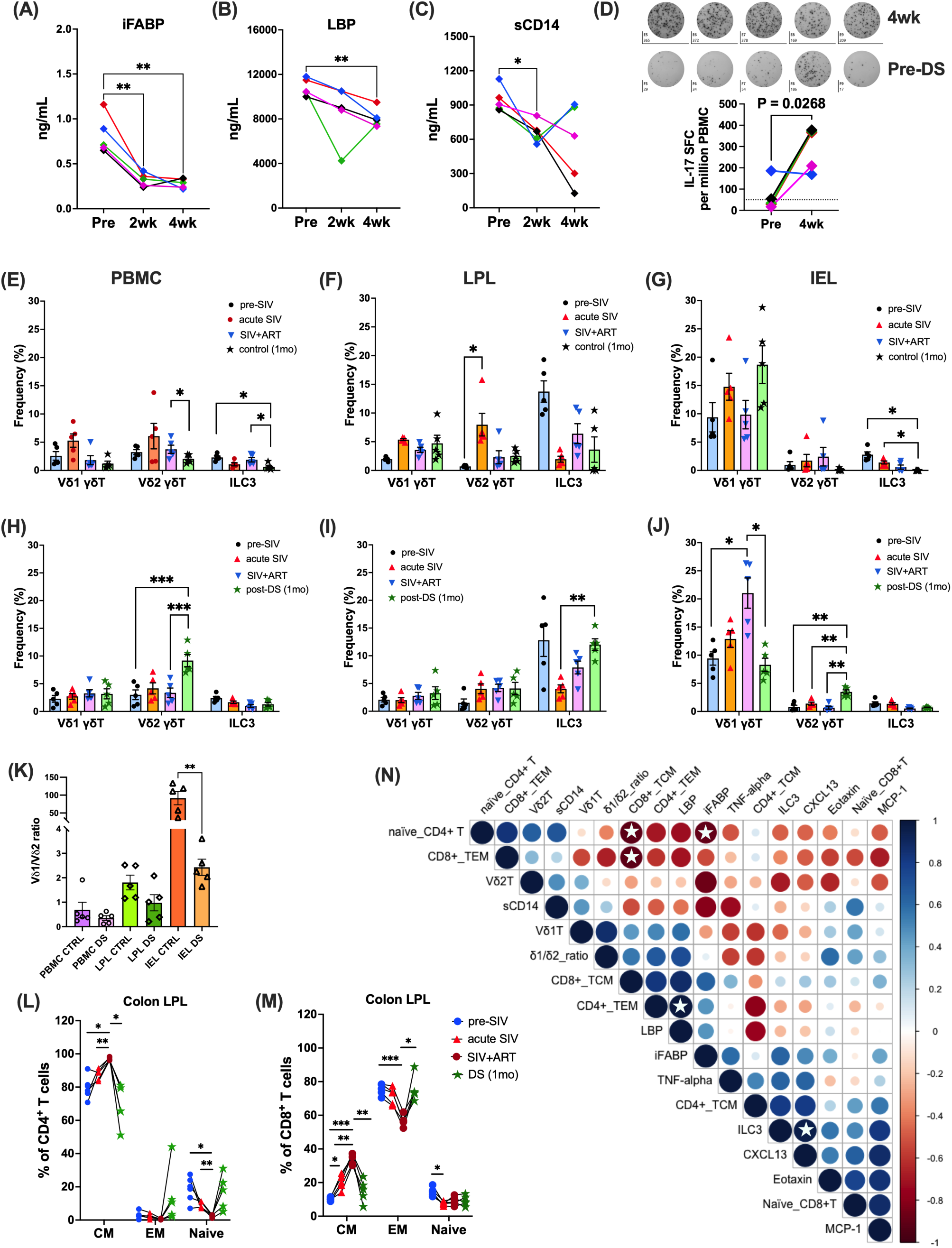
Effects of broccoli-based diet supplement on plasma biomarkers and immune cell frequencies in chronic SIV-infected macaques on long-term ART. Plasma iFABP **(A)**, LBP **(B)**, and sCD14 **(C)** at baseline, 2- and 4-weeks post-supplementation (950–1900 mg/day). **(D)** IL-17⁺ PBMC after Vδ2 T agonist stimulation at baseline and 4 weeks. **(E–J)** Frequencies of Vδ1 T, Vδ2 T, and ILC3 in PBMC, LPL, and IEL from control vs diet-supplemented (DS) groups. **(K)** Vδ1/Vδ2 ratios in PBMC, LPL, and IEL. **(L)** CD95⁺CD28⁺ central memory (CM) and CD95⁻CD28⁺ naïve CD4⁺ T cells; **(M)** CD95⁺CD28⁺ CM and CD95⁺CD28⁻ effector memory (EM) CD8⁺ T cells in colonic LPL after 4 weeks. Data shown as mean ± SEM; paired ANOVA (*p<0.05, **p<0.01, ***p<0.001). (N) Pearson correlations from −1 (red) to +1 (blue); white asterisks indicate significance.

Flow cytometry showed a significant increase in circulating Vδ2 T cells at 4 weeks in the DS group, contrasting with a decline in controls (Fig. 7E, H). In colonic LPL, DS increased ILC3 but did not alter γδ T cell frequencies (Fig. 7F, I). Intraepithelial lymphocytes exhibited reduced Vδ1 and increased Vδ2 frequencies (Fig. 7G, J), lowering the Vδ1/Vδ2 ratio across all compartments, most significantly in IEL (Fig. 7K). Classical T cell frequencies were unchanged (Supplementary Fig. 4), but DS reduced CD95⁺CD28⁺ central memory CD4⁺ T cells in LPL with a trend toward more naïve CD4⁺ T cells (Fig. 7L) and significantly decreased CD8⁺ central memory while increasing effector memory cells (Fig. 7M).

Correlation analysis linked immune subsets with plasma biomarkers (Fig. 7N). Naïve CD4⁺ T cells negatively correlated with iFABP consistent with their recovery post-DS, and ILC3 tracked with CXCL13 suggesting a gut mucosal–B cell chemokine axis. MCP-1, CXCL13, and Eotaxin clustered together, highlighting a shared inflammatory program spanning monocyte/macrophage activation, eosinophil recruitment, and lymphoid remodeling, which was dampened by DS (Supplementary Fig. 3). Overall, DS improved epithelial barrier markers, enhanced IL-17–producing Vδ2 responses, and reshaped mucosal T cell and ILC3 subsets during chronic SIV+ART.

In colonic LPL, DS significantly increased AhR in Vδ2 T cells (P = 0.01) with a trend in Vδ1 T cells (P = 0.06) but decreased AhR in ILC3 (P = 0.006) (Fig. 8A). RORγt increased across all subsets, most prominently in ILC3 (P = 0.003) and Vδ1 T cells (P = 0.04), while T-bet declined in Vδ1 T cells (P = 0.004) (Fig. 8B-C). These shifts indicate a move toward a Type 3 immune phenotype, supported by polyfunctional transcription factor expression profiles in Vδ1 T cells and ILC3 (Fig. 8D). Ex vivo stimulation showed increased IL-22 response in Vδ1 (P = 0.0008) and Vδ2 (P = 0.007), reduced IFN-γ, and unchanged IL-17A (Fig. 8E–G), with no significant changes in polyfunctional cytokine profiles (Fig. 8H-J). ILC3 cytokines were unaffected despite significantly increased RORγt expression. CD4⁺ and CD8⁺ T cells exhibited no functional recovery following supplementation. Their IL-17 and IL-22 responses continued to decline under chronic SIV+ART, consistent with progressive loss of epithelial barrier–protective functions and were accompanied by a marked reduction in IFN-γ production (Supplementary Fig. 5), indicative of dampened inflammatory capacity. Taken together, these findings demonstrate that dietary supplementation reprograms colonic γδ T cells toward Type 3 immunity characterized by increased IL-22 production, a shift likely to enhance mucosal barrier integrity and epithelial repair.

**Fig. 8.**
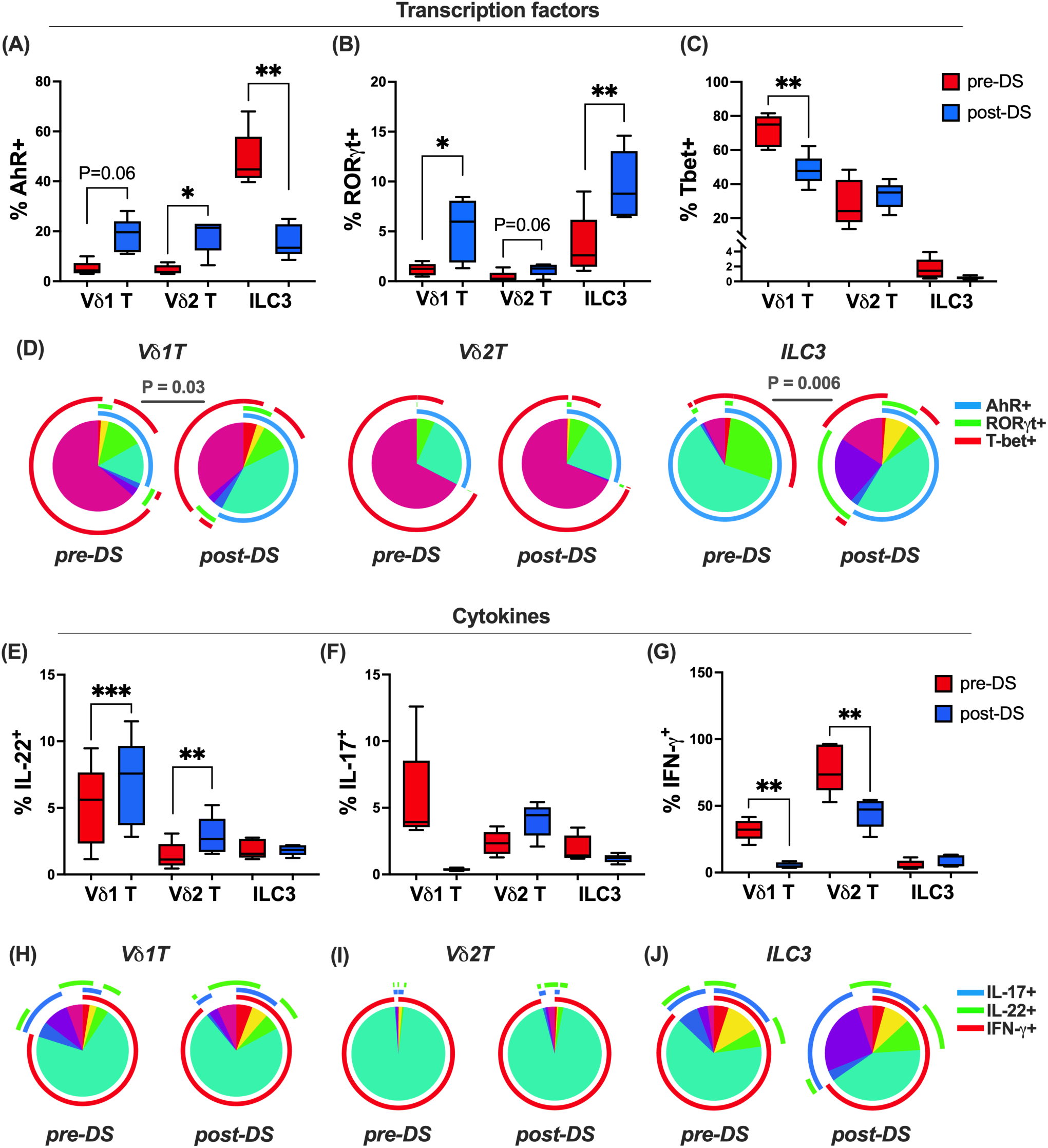
Effects of broccoli-based diet supplement on transcriptional immunophenotype and cytokine producing functions of colonic mucosal γδ T cells and ILC3 during chronic SIV+ART. Colonic lamina propria lymphocytes were analyzed for transcription factor expression and cytokine production following PMA/ionomycin stimulation. Frequencies of AhR **(A)**, RORγT **(B)**, and T-bet **(C)** expression were compared pre- and 1-month post-dietary supplementation (DS) with indoles. **(D)** Pie charts depicting changes in the relative expression of AhR, RORγT, and T-bet by Vδ1 T, Vδ2 T, and ILC3 populations in colonic LPL fraction from pre- and post-DS. Pie arcs represent expression of individual transcription factors and pie slices represent the number of co-expressed transcription factors. Cytokine production of IL-22 **(E)**, IL-17A **(F)**, and IFN-γ **(G)** in the same subsets was measured following mitogen stimulation of colonic LPL, and combinatorial cytokine responses shown in pie charts from pre- and post-DS **(H–J)**. Mean ± SEM; paired ANOVA (*p<0.05, **p<0.01, ***p<0.001, ****p<0.0001).

## Discussion

Despite effective ART, intestinal barrier dysfunction persists in people with HIV, underscoring gaps in our understanding of the mechanisms that sustain mucosal disruption. Our findings suggest that SIV infection and ART differentially shape epithelial integrity and mucosal immune regulation, with persistent defects in type 3 immunity pointing to AhR-dependent pathways as potential contributors to barrier homeostasis. Acute SIV infection was characterized by loss of ILC3 and Vδ2 T cells, along with compartment-specific alterations in the colon that reflected a localized Th1/Th17 imbalance and impaired epithelial–immune crosstalk. While ART partially reduced circulating markers of epithelial barrier disruption, it did not reverse SIV-driven Th1 polarization or restore type 3 immunity in the gut. Notably, AhR ligands attenuated SIV-induced epithelial injury in vitro, and dietary supplementation was associated with reduced barrier disruption and enhanced gut IL-22 responses, suggesting a mechanistic link between AhR signaling and the preservation of mucosal integrity during treated infection.

Our findings highlight that disruption of type 3 immunity is a central barrier to mucosal repair in HIV/SIV infection. While loss of CD4 T cells has long been recognized as a hallmark of disease, consistent with prior reports, our data point to a broader defect involving γδ T cells and ILC3s, populations critical for sustaining epithelial barrier integrity. The transient expansion and gut trafficking of Vδ2 T cells during acute infection suggest an early compensatory response, but one that is neither durable nor sufficient under ART. This dynamic may explain why epithelial tight junction restoration remains incomplete, as reflected by persistent elevations in zonulin despite reduced iFABP levels^25–27^. These results align with growing evidence that innate lymphocyte dysfunction underlies the failure of ART to fully normalize gut barrier function and inflammation^7, 14, 28^. Importantly, they suggest that interventions aimed at preserving or restoring type 3 immunity, whether by supporting γδ T cell and ILC3 function or by modulating barrier regulatory pathways, may be essential to achieve durable mucosal immune homeostasis. More broadly, our study reinforces the concept that immune restoration in HIV/SIV is not simply a matter of CD4 T cell recovery but requires a coordinated reconstitution of multiple mucosal immune cell networks that protect and repair the epithelial barrier.

In SIV-naïve macaques, colonic immunity is dominated by AhR⁺/RORγt⁺ γδ T cells and ILC3s, consistent with their roles in mucosal defense and tissue repair. However, following SIV infection, impaired IL-17A and IL-22 production by these cells correlated with reduced RORγt and AhR expression, revealing a mechanism by which SIV, despite ART, compromises gut barrier function. Our data further reveal that chronic SIV+ART reprograms γδ T cells and ILC3s toward a T-bet–driven, IFN-γ–biased phenotype, at the expense of IL-17 and IL-22. Although elevated T-bet supports HIV-specific CD8⁺ T cell cytotoxicity^29^ and may promote IL-22–producing ILCs^30^, production of IL-22 and IL-17 by intestinal immune cells is essential for epithelial integrity, antimicrobial defense, and microbiome homeostasis^31, 32^, and their loss likely sustains barrier dysfunction and microbial translocation during chronic HIV/SIV infections. The reciprocal regulation of IL-22 by RORγt and T-bet, and of IL-17 by AhR and T-bet, underscores the transcriptional plasticity of γδ T cells and ILC3s in mucosal inflammation^33, 34^. Notably, intraepithelial ILC3s remain persistently depleted despite ART, indicating durable disruption of epithelial immune integrity. Together, these findings suggest that transcriptional reprogramming of mucosal type 3 effector lymphocytes is a critical barrier to gut barrier restoration and identify RORγt and AhR signaling as potential therapeutic targets to re-establish mucosal immunity in treated HIV infection.

HIV-associated dysbiosis perturbs tryptophan metabolism and reduces microbiota-derived indoles, linking microbial imbalance to impaired type 3 immunity^6, 35^. Interventions aimed at restoring mucosal function have demonstrated that probiotics improved CD4⁺ T cell recovery and reduced inflammation in SIV-infected macaques, particularly with IL-21 supplementation^36, 37^, while fecal microbiota transplantation lowered gut permeability in ART-suppressed people with HIV^38^. Diet quality also influences mucosal immunity, with Mediterranean diet improving immune activation and microbiota composition^39^, whereas high-fat diet accelerated disease progression and microbial translocation^40^.

Because chronic HIV/SIV alters the microbiome and may impair microbial conversion of glucobrassicin into the AhR ligand I3C, we used a broccoli-based supplement with active myrosinase to release I3C independent of microbial metabolism. This intervention enhanced mucosal barrier integrity during treated SIV infection by restoring γδ T cell and ILC3 function, increasing naïve CD4⁺ and effector-memory CD8⁺ T cells in the colon, and lowering Vδ1:Vδ2 ratios in blood and intestinal compartments. Vδ1:Vδ2 inversion in both peripheral blood^41, 42^ and the gastrointestinal tract^43^ is a hallmark of HIV infection, which is driven in part by loss of gut mucosal Vδ2 cells and expansion of Vδ1 T cells. This inversion correlates with elevated intestinal barrier dysfunction biomarkers and systemic inflammation during long-term ART^9^. Importantly, similar Vδ1:Vδ2 imbalances are observed in other chronic inflammatory conditions affecting the gut, such as kidney disease, viral hepatitis, and obesity^8^, highlighting a broader link between γδ T-cell subset dysregulation, gut permeability, and chronic inflammation. Thus, the observed rebalancing in the DS group in our study suggests a mechanism for barrier protection, consistent with murine studies where I3C activated AhR, induced its target gene, Cyp1a1, restored intestinal γδ T cells^44^, and promoted IL-22-mediated epithelial repair^18^, effects abrogated by IL-22 blockade^45^. Notably, IL-22 upregulation occurred with I3C and not with butyrate, indicating mechanisms beyond butyrate production, since butyrate alone did not induce IL-22 expression. Consistently, butyrate supplementation in SIV-infected macaques failed to reduce microbial translocation or inflammation^46^, likely due to its predominant GPR-dependent rather than AhR-mediated signaling. These results establish a translational bridge between murine and primate gut immunity, highlighting the central role of the gut AhR pathway in SIV-induced barrier disruption.

Our findings indicated that dietary indole supplementation was associated with a trend of reduction across several classes of circulating inflammatory mediators, specifically lowering key cytokines, chemokines, and angiogenic factors like TNF-α, MCP-1, CXCL13, and VEGF-D. This suggests the supplements dampened immune activation and tissue remodeling pathways. Correlation analyses further revealed that the supplements attenuated a chemokine-driven inflammatory axis (MCP-1, CXCL13, eotaxin) while promoting gut barrier repair and mucosal immune balance. We also observed a restoration of naïve CD4⁺ T cells, and an association between ILC3 and CXCL13, pointing to integrated T cell and innate lymphoid contributions to gut immune remodeling. Though not statistically significant in this small cohort, trends toward reduced IL-1RA and SDF-1α suggest a decreased need for compensatory anti-inflammatory responses.

In conclusion, our study provides proof of principle that dietary indoles can modulate gut mucosal immunity during SIV-induced barrier disruption. While I3C is likely the primary bioactive compound, other metabolites such as sulforaphane may contribute, and microbiome effects were not prospectively assessed. Nonetheless, these findings highlight the role of AhR–IL-22 signaling in maintaining and repairing the intestinal barrier under ART-suppressed infection. The observed associations between type 3 immunity, AhR–RORγt signaling, and epithelial function generate testable hypotheses, and future studies using spatial transcriptional profiling will be critical to define how microbial- and diet-derived ligands modulate mucosal repair and barrier homeostasis.

## Methods

### Animals, Viral Inoculation, and ART

The animal study was reviewed and approved by Tulane University Institutional Animal Care and Use Committee (IACUC). Eleven healthy adult Indian-origin rhesus macaques (Macaca mulatta), aged 5–10 years and seronegative for SIV, HIV-2, STLV-1, SRV-1, and herpes-B, were assigned to control (n = 6) or diet supplement (DS, n = 5) groups based on prior SIV+ART studies. Animals were infected intrarectally with 2500 TCID50 SIVmac251 (Preclinical Research and Development Branch, NIAID). ART was administered daily via subcutaneous injection: 5.1 mg/kg Tenofovir Disoproxil Fumarate (TDF), 30 mg/kg Emtricitabine (FTC), and 2.5 mg/kg Dolutegravir (DTG) in 15% kleptose solution at pH 4.2^9^. Plasma viral loads were quantified using the Roche High Pure Viral RNA Kit^47^.

### Cell Isolation

Blood collected in EDTA tubes (Sarstedt) was processed immediately. PBMCs were isolated by density gradient centrifugation (Lymphocyte Separation Medium, MP Biomedicals) at 1500 rpm for 45 min for phenotyping and functional assays. Colonic intraepithelial and lamina propria lymphocytes were isolated as described^48^. Briefly, biopsies were washed with PBS, epithelial cells removed with RPMI-1640 + 5% FBS and 5 mM EDTA at 37°C for 1 h, and tissue digested with RPMI-5 + 60 U/ml Type II collagenase. LPLs were enriched via Percoll density gradients, washed, and resuspended in RPMI-10 + 10% FCS. Cell viability was >90% by trypan blue exclusion.

### Immunophenotyping

Polychromatic flow cytometry was performed using anti-human mAbs cross-reactive with rhesus macaques^9^. Fresh or frozen PBMCs (1–2 million) were stained with viability dye, surface markers (Supplementary Table S1), and transcription factors RoRγt, T-bet, and AhR following fixation (True Nuclear Kit, Biolegend). Unstained controls were included with each batch. Cells were resuspended in PBS + 1% formaldehyde and ≥200,000 lymphocytes were acquired on a BD Symphony; analysis was performed using FlowJo v10.

### Plasma Markers of Inflammation, Microbial Translocation, and Intestinal Damage

Frozen plasma was thawed, filtered (Ultrafree, Millipore), and used for multiplex quantification of 37 cytokines, chemokines, and growth factors (ProcartaPlex, Invitrogen) per manufacturer’s instructions. Data were acquired on a Bio-Plex 200 and analyzed with Bio-Plex Manager v6.1. Plasma IFABP, LBP, sCD14, and zonulin were measured using commercial ELISA kits (MyBioSource, R&D Systems, Alpco Diagnostics) in duplicate, and data were analyzed with Gen5 software (BioTek).

### Functional analyses

PBMCs and rectal lamina propria lymphocytes were resuspended at 1 × 10⁶ cells/mL in RPMI + 10% FBS and antibiotics, then stimulated 16 h at 37°C with PMA/ionomycin ± brefeldin A. Cells were stained for surface (CD45, CD3, CD4, CD8, TCR γδ/Vδ1/Vδ2, CD161) and intracellular markers (CD69, IL-17, IL-22, TNF-α, IFN-γ), fixed, and acquired on BD Symphony/LSRFortessa; analysis was via FlowJo v10. IL-17–secreting cells were measured by ELISPOT using PMA/ionomycin or HMBPP for γδ T cells; responses >2-fold above background and >50 SFC/10⁶ PBMC were considered positive.

### Effect of HIV-tat on Caco-2 Epithelial Barrier Integrity Monitored by xCELLigence RTCA

Caco-2 cells were seeded at 20,000 cells/well in DMEM with 20% FBS, 1% non-essential amino acids, and antibiotics. Cell adhesion was monitored in real time using impedance-based RTCA, with Cell Index (CI)^49^ normalized to pre-treatment values. After ∼48 h, cells were treated with medium, 3 μM SIVmac_239_ Tat (ARP-12765, NIH HIV Reagent Program), ± I3C (1 μg/mL) or FICZ (1 μg/mL) (Sigma-Aldrich product #17256, SML1489), and CI measured every 5 min in quadruplicate. For ZO-1 immunofluorescence, parallel monolayers in glass-bottom plates were treated 5 h, washed, fixed with 2% PFA, permeabilized with 0.2% Triton X-100, and blocked with 5% goat serum. Cells were incubated with anti-ZO-1 primary antibody (1:100) overnight, followed by Alexa Fluor 488 secondary (1:500) and DAPI nuclear stain. Imaging was performed on a Nikon Eclipse Ti2 microscope, and ZO-1 junctional integrity was qualitatively assessed. DR-EcoScreen cells were seeded (100,000/mL) in α-MEM + 5% FBS and antibiotics, incubated overnight, and treated with test samples for 24 h. Luminescence was measured after adding Steady-Glo™ reagent and shaking 5 min at room temperature^50^.

### Statistical analyses

As appropriate, unpaired or paired, two-way t-tests and two-way ANOVAs or mixed effects models were used in statistical analyses of plasma analyte concentrations, lymphocyte phenotype and function frequencies (Prism v9.0, GraphPad Software Inc.). Significance of polyfunctional cytokine expression was assessed using the Spice permutation-test on relative expression values. Correlations were calculated in R using the Hmisc package and visualized with corrplot.

## Acknowledgments

We thank the Tulane National Primate Research Center (RRID:SCR_008167), its veterinary staff, and the flow cytometry core (S10 OD026800, RRID: SCR_024611/ SCR_008167). We also acknowledge ViiV and Gilead for antiretroviral drugs, the NIH Nonhuman Primate Reagent Resource for anti-α4β7-PE antibody, and the NIH HIV Reagent Program (ARP-12765) for SIV Tat. Funders had no role in study design, data collection, analysis, or manuscript preparation.

## Author contributions

**Siva Thirugnanam:** Writing – review & editing, Writing – original draft, Methodology, Formal analysis, Data curation, Conceptualization. **Alison R Van Zandt:** Methodology, Investigation, Formal analysis. **Alexandra B McNally:** Writing – review & editing. **Victoria A Hart:** Methodology, Data curation. **Isabelle Berthelot:** Writing – review & editing. **Lara A Doyle-Meyers:** Investigation. **David A Welsh:** Writing – review & editing, Conceptualization. **Andrew G MacLean:** Writing – review & editing, Investigation. **Namita Rout:** Writing – review & editing, Supervision, Project administration, Funding acquisition, Conceptualization.

