## Supplemental Data for "Dietary Indoles Modulate Gut Barrier Integrity via the AhR–IL-22 Axis in ART-Treated SIV Infection"

Address: 18703, Three Rivers Road, Covington, LA 70433, USA.

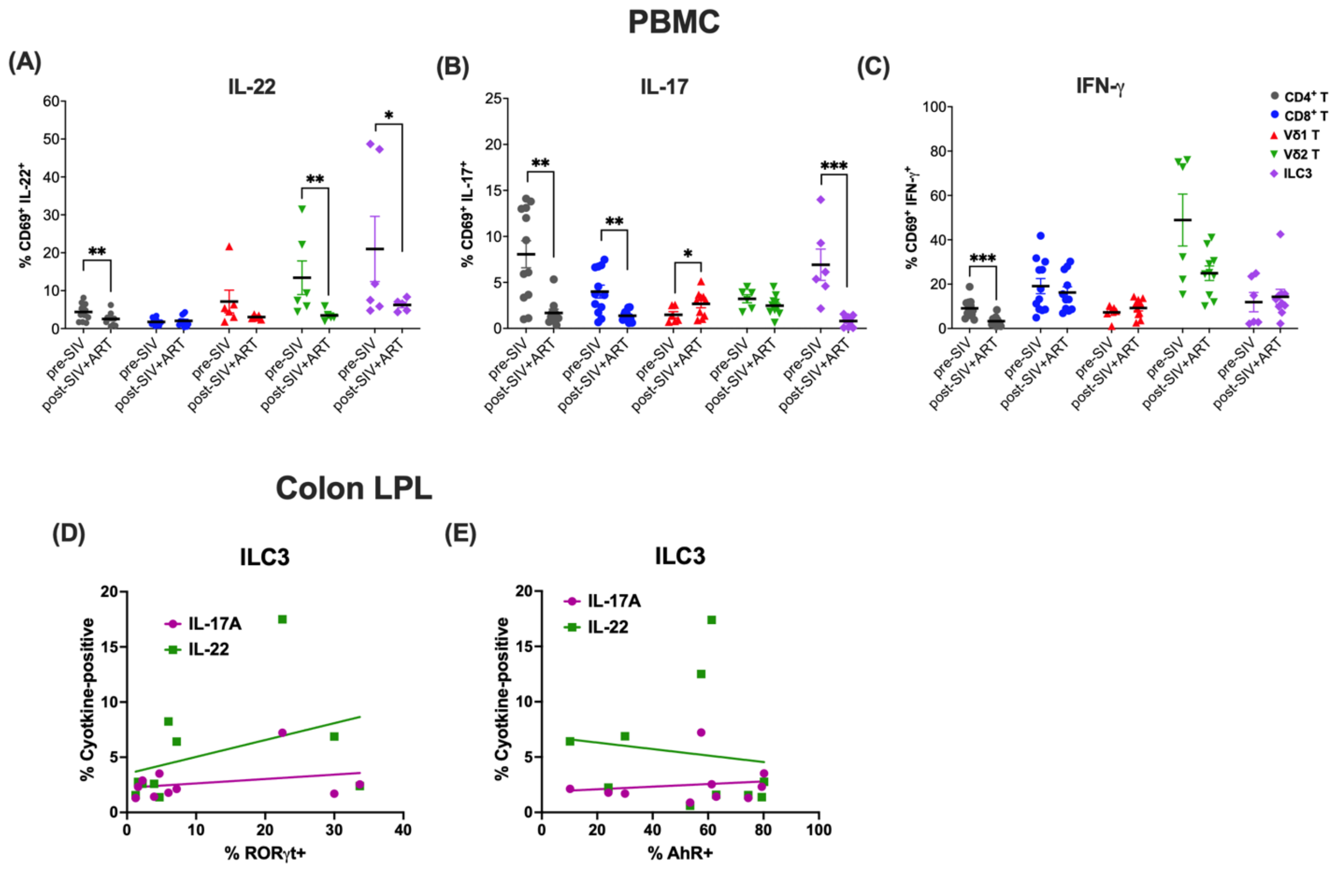


**Supplemental Figure 1. Cytokine production by peripheral T cells and ILC3 during ART-suppressed SIV infection and correlation of colonic LPL ILC3 cytokines with transcription factor expression.** Frequencies of $CD4^{+}$ T cells, $CD8^{+}$ T cells, γδ T cells, and ILC3s expressing IL-22 **(A)**, IL-17A **(B)**, and IFN-γ **(C)** determined by intracellular flow cytometry. Data represent mean ± SEM from 5-15 animals per group. Mean ± SEM; Mann-Whitney test (*p<0.05, **p<0.01, ***p<0.001). Spearman correlation analysis of RORγt **(D)**, and AhR **(E)** expression with IL-17A and IL-22 cytokines produced by colonic ILC3 populations.

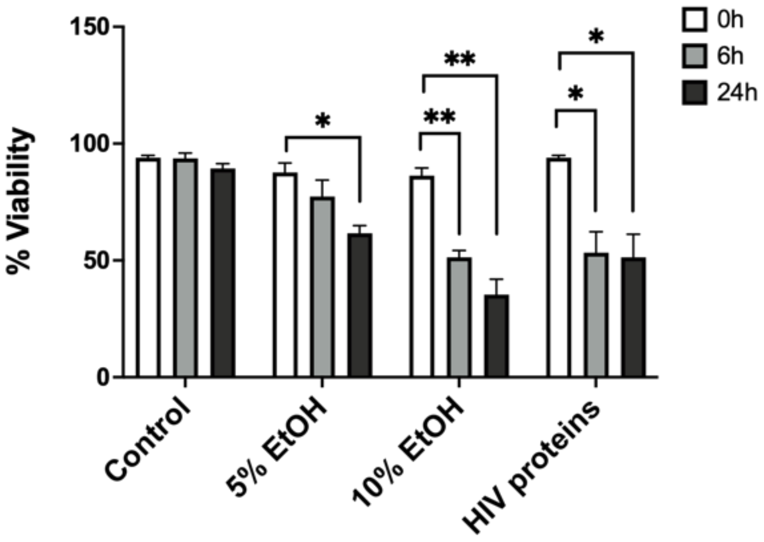


**Supplemental Figure 2. Temporal effects of HIV proteins and ethanol on CaCo-2 cell viability.** Caco-2 cell viability after the treatment with 5% EtOH, 10%EtOH, or HIV TAT+Nef+gp120 at 1μg/mL concentration. Viability was measured by trypan blue exclusion of dead cells at the indicated time-points. Paired t-test comparisons with matched wells at 0h prior to adding the treatments (*p<0.05, **p<0.01). Caco-2 cells alone served as negative controls at the indicated time-points.

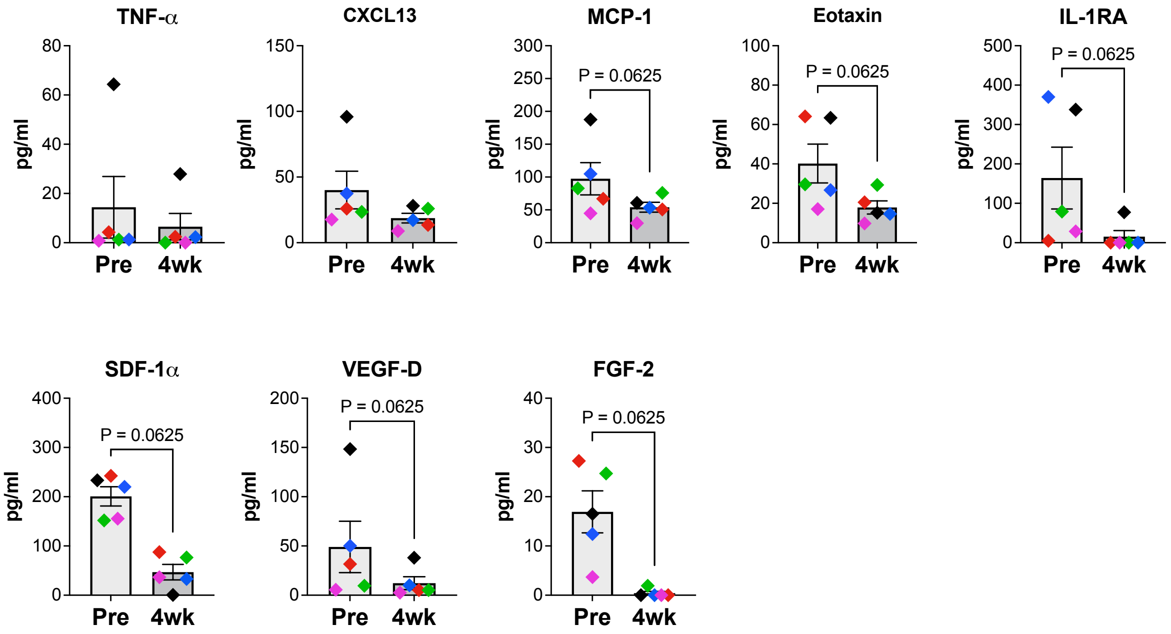


**Supplemental Figure 3. Effects of diet supplement on plasma levels of inflammatory cytokines and growth factors during SIV infection and ART.** Plasma levels of the inflammatory cytokines, chemokines, and growth factors: TNF-α, CXCL13, MCP-1, Eotaxin, IL-1RA, Stromal Cell-Derived Factor 1 alpha (SDF-1α), Vascular Endothelial Growth Factor D (VEGF-D), and Fibroblast Growth Factor 2 (FGF-2), evaluated at 6 month-post-SIV+ART (pre-DS) and 4-weeks post-DS.

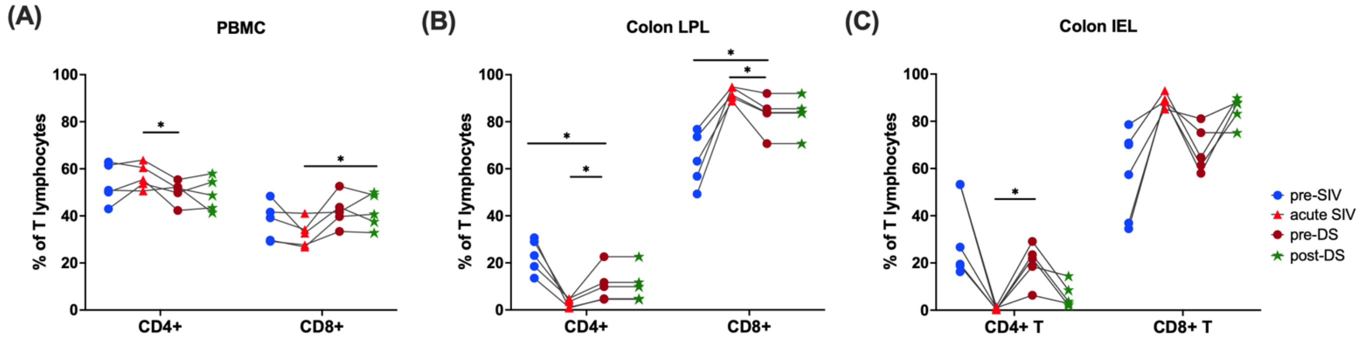


**Supplemental Figure 4. Frequencies of classical T cells during chronic SIV+ART and following 4 weeks of broccoli-based diet supplementation.** Ex vivo frequencies of $CD4^+$ T cells and $CD8^+$ T cells in **(A)** PBMC **(B)** colonic LPL, and **(C)** colonic IEL assessed longitudinally by flow cytometry. Data represent mean ± SEM in 5 animals: paired ANOVA (*$p < 0.05$).

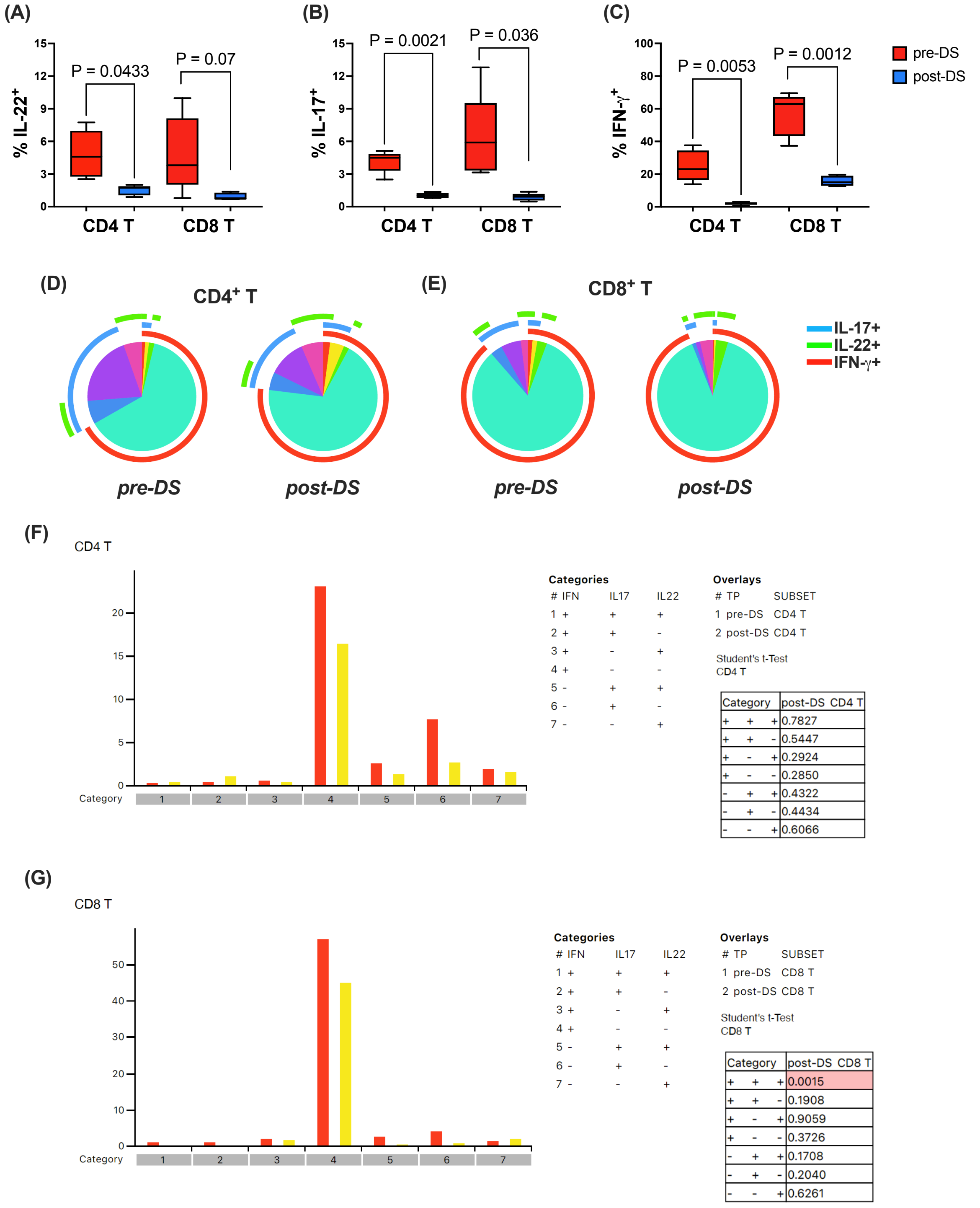


**Supplemental Figure 5. Cytokine producing functions of classical T cells in colonic LPL of rhesus macaques during chronic SIV+ART and following 4 weeks of broccoli-based diet supplementation.** Intracellular cytokine staining for IL-22 **(A)**, IL-17A **(B)**, and IFN-γ **(C)** in CD4+ T cells and CD8+ T cells following PMA/ionomycin stimulation of colonic LPL. Data represent mean ± SEM in 5 animals; paired ANOVA test. Pie charts depicting changes in the intracellular expression of cytokines by CD4+ T cells **(D)** and CD8+ T cells **(E)** in colonic LPL fraction from pre- and post-DS. Differences in each combination of IFN-γ, IL17A, and IL-22 producing cells at pre-DS (red bar) and post-DS (yellow bar) time-points in colonic CD4+ T cells **(F)** and CD8+ T cells **(G).**

**Table S1.** Phenotyping panel (Antibodies used for surface markers and transcription factor staining):

| Marker | Clone | Company | Fluorochrome |
|---|---|---|---|
| TCR Vδ1 | TS8.2 | Invitrogen | FITC |
| RoRγt | AFKJS-9 | Invitrogen | PCP-Cy5.5 |
| T-bet | 4B10 | Biolegend | BV421 |
| TCR γδ | B1 | Biolegend | BV510 |
| CD4 | OKT4 | Biolegend | BV605 |
| CD8 | SK1 | BD | BV650 |
| CD14 | M5E3 | BD | BV711 |
| CD45 | D058-1283 | BD | BUV395 |
| CD95 | DX2 | BD | BUV737 |
| CD20 | 2H7 | BD | BUV805 |
| a4b7 | A4B7R1 | NHP Resource | PE |
| AhR | T49-550 | BD | PE-CF594 |
| CD28 | CD28.2 | BD | PE-Cy5 |
| CD127 | eBioRDR5 | Invitrogen | PE-Cy5.5 |
| TCR Vδ2 | 15D | Thermoscientific | APC |
| FVS700 | ---- | BD | AL700 |
| CD3 | SP34-2 | BD | APC-Cy7 |

**Table S2.** Intracellular Cytokine Staining panel:

| Marker | Clone | Company | Fluorochrome |
|---|---|---|---|
| TCR Vδ1 | TS8.2 | Invitrogen | FITC |
| IL-22 | IL22JOP | Invitrogen | PCP-Cy5.5 |
| CD4 | OKT4 | Biolegend | BV605 |
| CD8 | SK1 | BD | BV650 |
| CD14 | M5E3 | BD | BV711 |
| CD45 | D058-1283 | BD | BUV395 |
| PD-1 | EH12.1 | BD | BUV615 |
| CD95 | DX2 | BD | BUV737 |
| CD20 | 2H7 | BD | BUV805 |
| IL-17A | eBio64DEC17 | Invitrogen | PE |
| CD69 | FN50 | BD | PE-CF594 |
| CD161 | DX12 | BD | PE-Cy5 |
| CD127 | eBioRDR5 | Invitrogen | PE-Cy5.5 |
| IFN-$\gamma$ | B27 | BD | PE-Cy7 |
| TCR Vδ2 | 15D | Thermoscientific | APC |
| FVS700 | --- | BD | AL700 |
| CD3 | SP34-2 | BD | APC-Cy7 |
